# Virus-inclusive single-nucleus RNA sequencing reveals two distinct endothelial response patterns in infectious salmon anaemia

**DOI:** 10.1101/2025.03.20.644265

**Authors:** Adriana Magalhães Santos Andresen, Richard S. Taylor, James J. Furniss, Maryam Saghafian, Raoul Valentin Kuiper, Daniel J. Macqueen, Johanna Hol Fosse

**Author notes:** Corresponding authors: (AMSA); (JHF).

## Abstract

Viral replication in endothelial cells is a hallmark of many viral diseases in humans and other animals, underscoring the importance of understanding cellular mechanisms that restrict viral replication and the associated consequences for vascular health. Pathogenic variants of infectious salmon anaemia virus (ISAV, *Isavirus salaris*) target endothelial cells of Atlantic salmon (*Salmo salar L.*), causing severe systemic disease and major losses during outbreaks in aquaculture. To better understand the endothelial response to ISAV, we used single nucleus RNA-sequencing at pre-clinical (12 days post infection, dpi) and clinical (16 dpi) stages of infection. Our approach enables an assessment of transcriptomic responses for different endothelial subpopulations at unprecedented resolution. ISAV RNA was predominantly detected in endothelial cells, which, along with mononuclear phagocytes, showed the highest number of differentially regulated genes at both time points. At 12 dpi, differentially expressed genes in endothelial cells were enriched for pathways related to NOD-like receptor signaling, antiviral responses, and regulation of programmed cell death. By 16 dpi, we observed a shift toward enrichment of pathways associated with cellular senescence, apelin signaling, and insulin signaling. We identified two distinct infection-related states at both time points: a virus-permissive state characterized by upregulation of genes involved in protein synthesis, small GTPase signaling, and MAPK activity, and a bystander phenotype marked by activation of antiviral responses, immune signaling, and translational regulation. This study is the first to capture the individual cell type responses to ISAV infection, and to characterize the in vivo endothelial response to active viral replication at single-cell resolution in any species.

## Introduction

Endothelial cells orchestrate immune and inflammatory responses and mitigate immune pathology during viral infection, and endothelial structural and functional damage is a common feature of severe viral conditions (1). Pathological changes can affect both infected and non-infected endothelium and may have severe consequences, including bleeding, vascular leakage, and disturbed coagulation. Importantly, affordable interventions that non-specifically stabilize endothelial function have shown promise in managing viral infections causing life-threating vascular damage (2). Infectious salmon anaemia virus (ISAV) is an orthomyxovirus that causes disease in farmed Atlantic salmon. It is a significant economic threat to the aquaculture industry, and pathogenic ISAV is notifiable to both the World Organisation for Animal Health and the European Union. ISAV outbreaks lead to severe economic losses associated with premature slaughter, mandatory site fallowing, potential trade restrictions, and the implementation of preventive measures such as vaccination (3). Between 2007 and 2009, an ISAV epidemic in Chile had an estimated direct economic impact of US$2 billion (4). More recently, in 2021, Norway recorded 25 ISAV outbreaks – the highest number reported since the early 1990s (3).

In individual fish, ISAV infection begins in the surface epithelium before spreading to vascular endothelial cells of inner organs (5), with release of new viral particles into the blood (6). Clinical signs of infectious salmon anaemia (ISA) include anaemia, bleeding, ascites, and circulatory failure, often resulting in death (7). The endotheliotropic nature of ISAV is interesting, because of the general need to explore the vascular response to viral infection and its relation to the progression of infection and disease outcomes. Several human viruses, such as henipaviruses and hantaviruses, specifically target and replicate in endothelial cells, while others infect these cells as part of a broader cellular tropism (1). Avian influenza in poultry and bluetongue in ruminants are examples of viral diseases in terrestrial food production animals with widespread infection of endothelium (8, 9).

Single cell RNA-seq (scRNA-seq) approaches have significantly advanced our understanding of the dynamic heterogeneity of vascular endothelial cells across organs and in various pathogenic conditions (10). Given that respiratory pathology has been a key feature of recent viral pandemics, most scRNA-seq studies of the human endothelial response to viral infection have focused on the pulmonary vasculature (11, 12). Notably, the endothelium does not appear to contribute to viral amplification of SARS-CoV-2 or influenza A viruses in humans (13, 14). Moreover, because analysis of the tissue-resident endothelial cells requires biopsies of infected organs, together with the severe nature of disease, investigations in humans are usually limited to terminal cases. We therefore need complementary studies that cover the active phase of viral replication and extra-pulmonary organs to deepen our understanding of the relationship between endothelial responses and viral replication, dissemination, and pathology.

The aim of this investigation was to map the Atlantic salmon response to ISAV infection using single nucleus RNA-seq (snRNA-seq) of an immune and haematopoietic organ (head kidney), providing a detailed examination of the endothelial response to virus infection in a natural host. Samples were collected from fish infected with the high-virulent isolate NO/Glesvaer/2/90 and non-infected controls at two time points, before and after the onset of clinical signs. At both times, viral RNA was predominantly detected in endothelial cells, which also showed a marked transcriptomic response to infection. Two distinct infection-related endothelial transcriptional states were identified, one bystander-like with high expression of protective antiviral genes and one with a high proportion of directly infected endothelial cells, characterized by expression of stress and inflammatory markers. Our study provides comprehensive new insights into cell type responses to ISAV infection, capturing the *in vivo* endothelial response to active viral replication for the first time in any species.

## Results

### Experimental design, viral dynamics, mortality, and tissue pathology

To resolve head kidney cell type responses to ISAV infection, we performed snRNA-seq using Atlantic salmon infected with the high-virulent Norwegian ISAV isolate NO/Glesvaer/2/90 (15, 16). Fig 1A shows the infection trial design and experimental setup. We have previously described the progression of anaemia, viremia, and disease in the same trial (17). Fig 1B summarizes several key parameters, including the temporal dynamics of viral RNA detection in head kidney and blood and the associated cumulative mortality. The analysed time points correspond to when ISAV RNA levels peaked in head kidney (12 dpi) and blood (16 dpi), respectively. During sample collection, external organs were inspected and pathological findings recorded. Fig 1C illustrates gross pathology consistent with ISA disease. At 12 dpi, gross organ morphology in infected fish appeared normal, except for an enlarged spleen. At 16 dpi, we observed typical ISA pathology, with a darkened and enlarged liver, ascites, and petechial hemorrhaging of intra-abdominal fat and intestines. The first death was detected 15 dpi, and the trial was terminated 18 dpi, at its predetermined humane endpoint (40% cumulative mortality).

**Fig 1.**
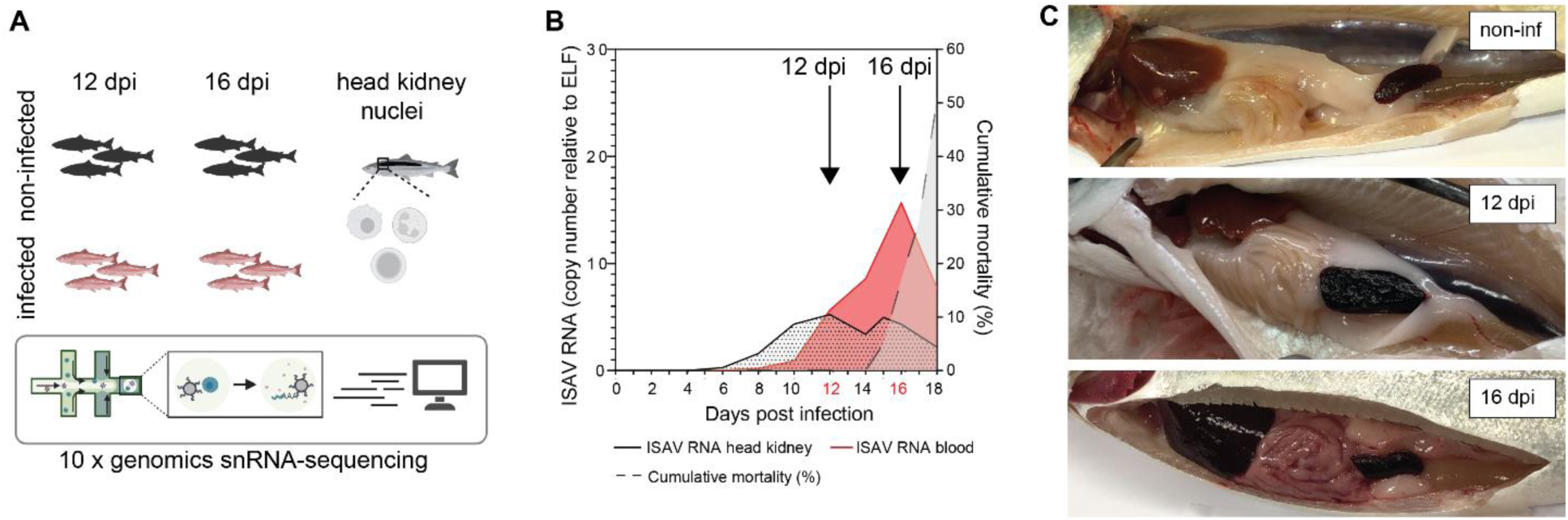
– Experimental infectious salmon anaemia. (A) Trial design and experimental setup. (B) RT-qPCR quantification of ISAV segment 8 RNA levels in head kidney and blood in relation to cumulative mortality. The same data have been presented in a previous publication (17), but is depicted here for context. (C) Gross appearance of abdominal organs of non-infected (non-inf) and virus-infected fish 12 and 16 dpi. Panel A was created with BioRender.com.

### Head kidney cell types in ISAV-infected and healthy Atlantic salmon

We recovered 61,112 nuclei from the 12 head kidney samples. After quality control, including filtration of poor quality nuclei and elimination of doublets, 56,421 nuclei remained for downstream analysis. Using an automated annotation pipeline (18), we identified 11 distinct major cell populations: B cells, endothelial cells, erythrocytes, granulocytes, hematopoietic stem cells (HSC), interrenal-like cells, mesenchymal cells, mononuclear phagocytes (MPs), natural killer-like cells (NK-L), T cells, and thrombocytes (Fig 2A). S1 Table displays the list of all marker genes for each cell population. Most cell populations were retrieved from all sample groups, except for natural killer-like cells that were not retrieved from infected fish at 12 dpi. S1 Fig show the expression levels of known marker genes on each cell population and individual variation of the distribution of cell types.

**Fig 2.**
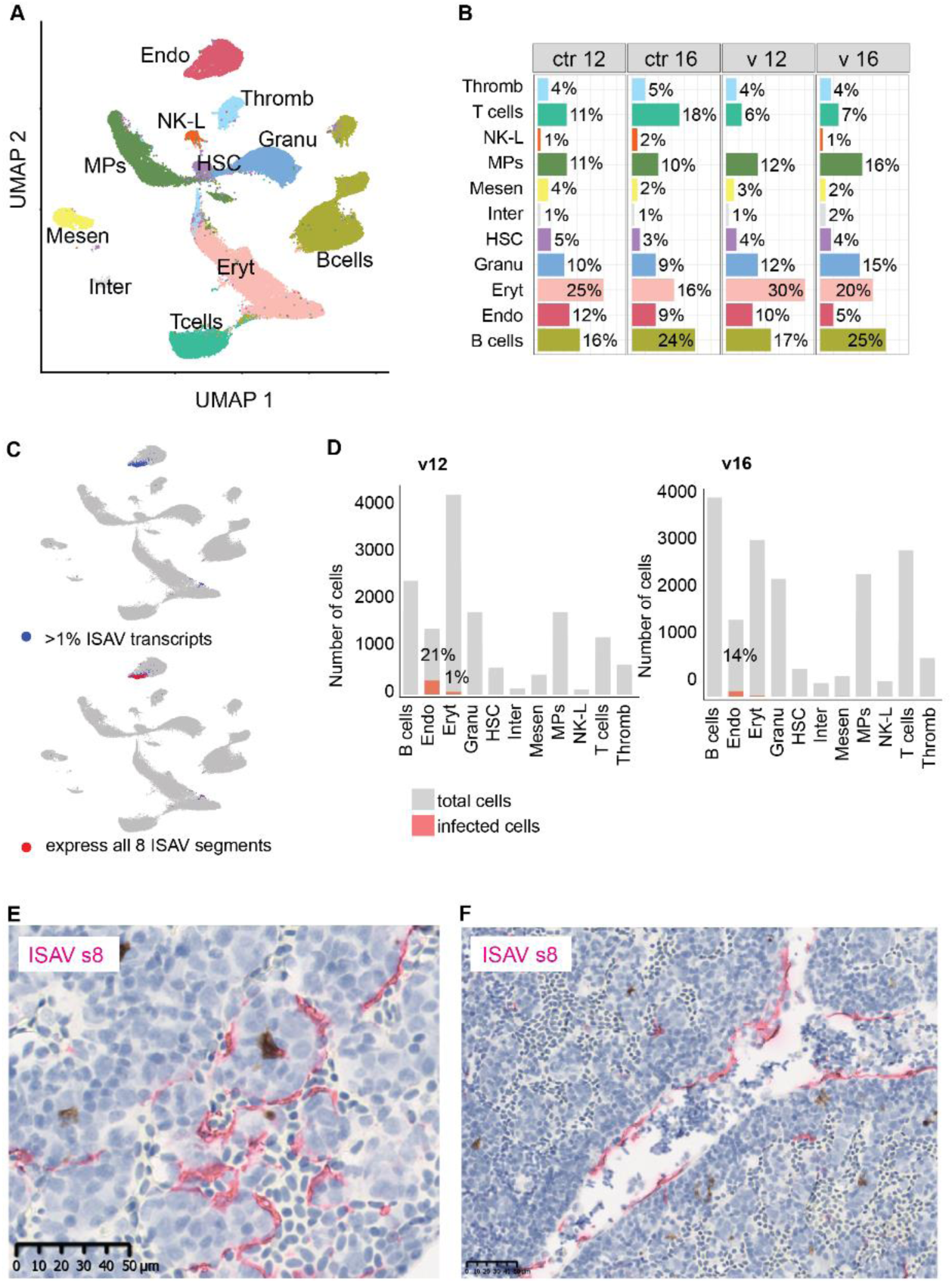
Head kidney cell types and distribution of ISAV RNA. (A) UMAP of merged samples showing the major cell populations identified in Atlantic salmon head kidney. (B) Barplot showing the percentage of cells in each cluster for each sample group: non-infected fish 12 dpi (ctrl12) and 16 dpi (ctrl16), and infected fish 12 dpi (v12) and 16 dpi (v16). (C) Feature plot showing ISAV gene segment RNA expression across all cell groups and conditions (control and virus-infected, both time points). Cells in blue are considered infected based on at least 1% of transcripts originating from ISAV RNA. Cells shown in red express all eight ISAV gene segments. Cells in gray do not express ISAV RNA. (D) Percentage of infected cells in each cell type in infected fish at 12 dpi (v12) and 16 dpi (v16). Abbreviations as follow: Thromb – thrombocytes, NK-L – natural killer-like cells, MPs – mononuclear phagocytes, Mesen – mesenchymal cells, Inter – interrenal cells, HSCs – haematopoietic stem cells, Granu – granulocytes, Eryt – erythrocytes, and Endo – endothelial cells. RNAscope in situ hybridization of ISAV segment 8 RNA (ISAV s8) in sections of head kidney from infected Atlantic salmon (12 dpi). Positive signal was observed in endothelial cells, both in thin-walled sinusoidal capillaries (E) and large vessels (F). Values for percentage of cells and infected cell in individual fish are provided in S1 Fig.

### The endothelium is the primary site of productive ISAV infection

To complement current knowledge of ISAV tropism (5, 19, 20), we next examined the expression of ISAV RNA across individual cells in our snRNA-seq dataset. This was possible because ISAV RNA is transcribed in the host cell nucleus and polyadenylated, which allows its detection by oligo-dT priming using the 10x Genomics platform. Since ambient viral RNA can make it challenging to determine which cells are infected, we set a threshold to differentiate between infected and non-infected cells, considering a cell as infected if at least 1% of its transcripts originated from viral RNA. A feature plot visualizing the combined expression of ISAV gene segments in both control and virus-infected samples identified 471 infected cells (Fig 2C, blue). ISAV RNA was primarily detected in endothelial cells, confirming its expected endothelial tropism. The same pattern was seen when visualizing cells that expressed all eight viral segments simultaneously (Fig 2C, red). Again, the majority of these cells (93%; 233 out of 250) were endothelial cells, reinforcing their critical role in ISAV infection. The number of cells infected and expressing each populations, with 21% of endothelial cells infected at 12 dpi and 14% at 16 dpi. While some erythrocytes also expressed ISAV genes, the fraction of positive cells was low (1% at 12 dpi). S1 Fig show the individual variation in the proportion of infected cells in endothelial and erythrocyte populations. In situ hybridization revealed ISAV RNA expression in endothelial cells, such as those lining thin-walled sinusoids and larger vessels in head kidney (Fig 2 E-F, S2 Fig). The same strict endothelial tropism was observed in heart and liver (S2 Fig).

### Endothelial cells mount a strong response to ISAV infection

To resolve cell type responses to ISAV infection, we identified the differentially expressed genes (DEGs) in each cell population using a pseudobulk approach, which aggregates counts from individual cells to enhance statistical power while accounting for biological variability across replicates, reducing false positives (21, 22). Fig 3 displays the number of DEGs for each cell type. The threshold for differential expression was set to adjusted p-value (padj) < 0.05 and log2 fold change (FC) > 1 for upregulated genes and < –1 for downregulated genes. MPs and endothelial cells mounted the strongest transcriptional response to ISAV infection, with the highest numbers of both upregulated and downregulated genes at both time points (Fig 3A). The full list of DEGs for each cell type is provided in S3 Table. In all cell types, a considerable proportion of upregulated DEGs overlapped with a recently characterized list of conserved antiviral response genes (23) (Fig 3B), cell types with a larger overlap may exhibit a stronger antiviral response.

**Fig 3.**
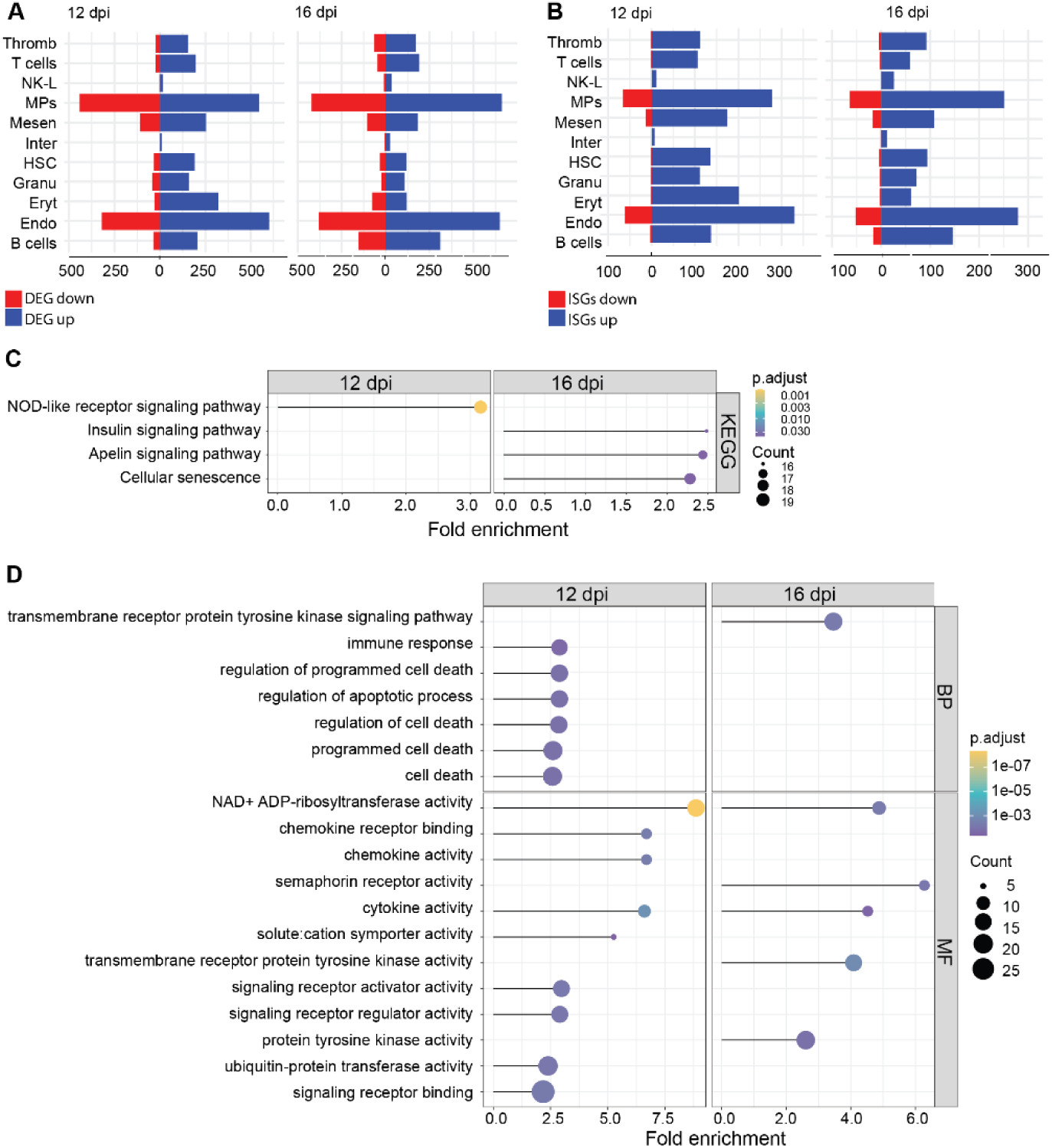
Endothelial cells mount a strong transcriptomic response to ISAV infection. (A – B) Bidirectional bar charts illustrate upregulated (blue) and downregulated (red) genes in each cell type. (A) Number of differentially expressed genes (DEGs) at 12 dpi and 16 dpi. (B) Number of differentially expressed interferon stimulated genes (ISGs) (23) at 12 dpi and 16 dpi. (C – D) Enrichment analyses of the endothelial response (combined up and downregulated) with a padj cutoff of 0.05. (C) KEGG enrichment analysis. Endo12 and Endo16 represent endothelial cells from 12 and 16 dpi, respectively. (D) Gene Ontology terms associated with biological processes (BP) and molecular functions (MF). In the dotplot, the lines represent fold enrichment, the colours indicate the padj and the dot sizes reflect the gene count.

### Endothelial transcriptomic response to ISAV

To decipher the overall response pattern of endothelial cells, we performed a gene set functional enrichment analysis using clusterProfiler (24). For each cell population, we generated one list containing all DEGs from pseudobulk analysis, including both upregulated and downregulated genes, and assessed the enrichment of gene ontology (GO) terms and KEGG pathways, applying a padj cutoff of 0.05. A list of enriched terms from the different groups is provided in S4 Table.

The endothelial transcriptomic response at 12 dpi was consistent with pathogen recognition and pro-inflammatory activation, with enrichment for the KEGG term NOD-like receptor signaling pathway (Fig 3C). Additionally, enriched GO terms indicated changes in immune response, regulation of programmed cell death, NAD+ADP-ribosyltransferase activity, ubiquitin protein transferase activity, cytokine and chemokine activity, and signaling receptor binding (Fig 3D).

At 16 dpi, there was a shift in the endothelial response, with enrichment of the KEGG pathways apelin signaling, insulin signaling, and cellular senescence (Fig 3C). There was also enrichment for transmembrane receptor protein tyrosine kinase signaling, as well as terms indicative of stress and immune responses, such as NAD+ADP-ribosyltransferase activity and cytokine activity (Fig 3D).

### Subclustering reveals bystander and directly infected endothelial cells

As endothelial cells are the primary target of ISAV infection and are also highly heterogeneous (5, 10), we next explored the heterogeneity of the endothelial response to ISAV. The 4,941 endothelial nuclei formed six distinct subpopulations (E1-E6) (Fig 4A). Marker genes of the different subpopulations were identified, including cells from both non-infected and infected fish at both time points (S5 Table). E1 was the most abundant subpopulation in non-infected fish, comprising 64% and 70% of all endothelial cells at 12 dpi and 16 dpi, respectively (Fig 4B). In infected fish, E1 diminished markedly to 10% at 12 dpi and 21% at 16 dpi (Fig 4B). The E1 transcriptional profile was consistent with scavenger-like capillaries (Fig 4C), the most abundant cell population in the teleost head kidney (18, 25, 26). In line with our previous findings in healthy fish (18), scavenger-like endothelial cells showed upregulation of *flt4* (ENSSSAG00000084309), *stab2* (ENSSSAG00000045007), *mrc1*(ENSSSAG00000073565), *meis2* (ENSSSAG00000007719) and *maf* (ENSSSAG00000072045), the latter two genes encoding transcription factors that induce markers of liver sinusoid endothelial cells (LSEC) (27). Overall, there was a high correlation (R = 0.91) between E1 marker genes in our current study and previously identified markers of head kidney scavenger-like endothelial cells (18) (Fig 4D). E2 was the second-largest subpopulation in our study, containing 28% and 21% of endothelial cells in non-infected fish and 39% and 27% of endothelial cells in infected fish at 12 and 16 dpi, respectively (Fig 4B). E6, representing ∼1% of endothelial cells, showed a very similar transcriptome profile to E2 (Fig 4C-D). Markers of E2 and E6 showed the closest correlation (R = 0.87 and R = 0.76, respectively) with markers of a subpopulation previously described in healthy Atlantic salmon head kidney (18) (Fig 4C-D), with enrichment for the transcription factors genes *ebf1* (ENSSSAG00000070298), *bach2* (ENSSSAG00000042426), and *runx1* (ENSSSAG00000097353), as well as genes involved in small G protein signaling, including *arhgap15* (ENSSSAG00000081610), *ahrgap24* (ENSSSAG00000044697), *ahrgap42b* (ENSSSAG00000105303), and *grb2* (ENSSSAG00000000363). The identity of this cluster remains undetermined. E4 contained 6% and 7% of endothelial cells in non-infected fish at 12 dpi and 16 dpi, and 3% of endothelial cells in infected fish at both time points (Fig 4B). E4 markers correlated most closely with previously identified markers of arterial (R = 0.90) and venous (R = 0.81) large vessel endothelial cells in the head kidney (18). While cells in E4 consistently presented as one cluster in our analysis, the distribution of markers was heterogeneous, with arterial markers such as *mecom* (ENSSSAG00000048049, ENSSSAG00000041461) and *gja5* (ENSSSAG00000042461) enriched in one region, while venous markers such as *vwf* (ENSSSAG00000101168) and *lama5* (ENSSSAG00000108070) were enriched in another (S3 Fig).

**Fig 4.**
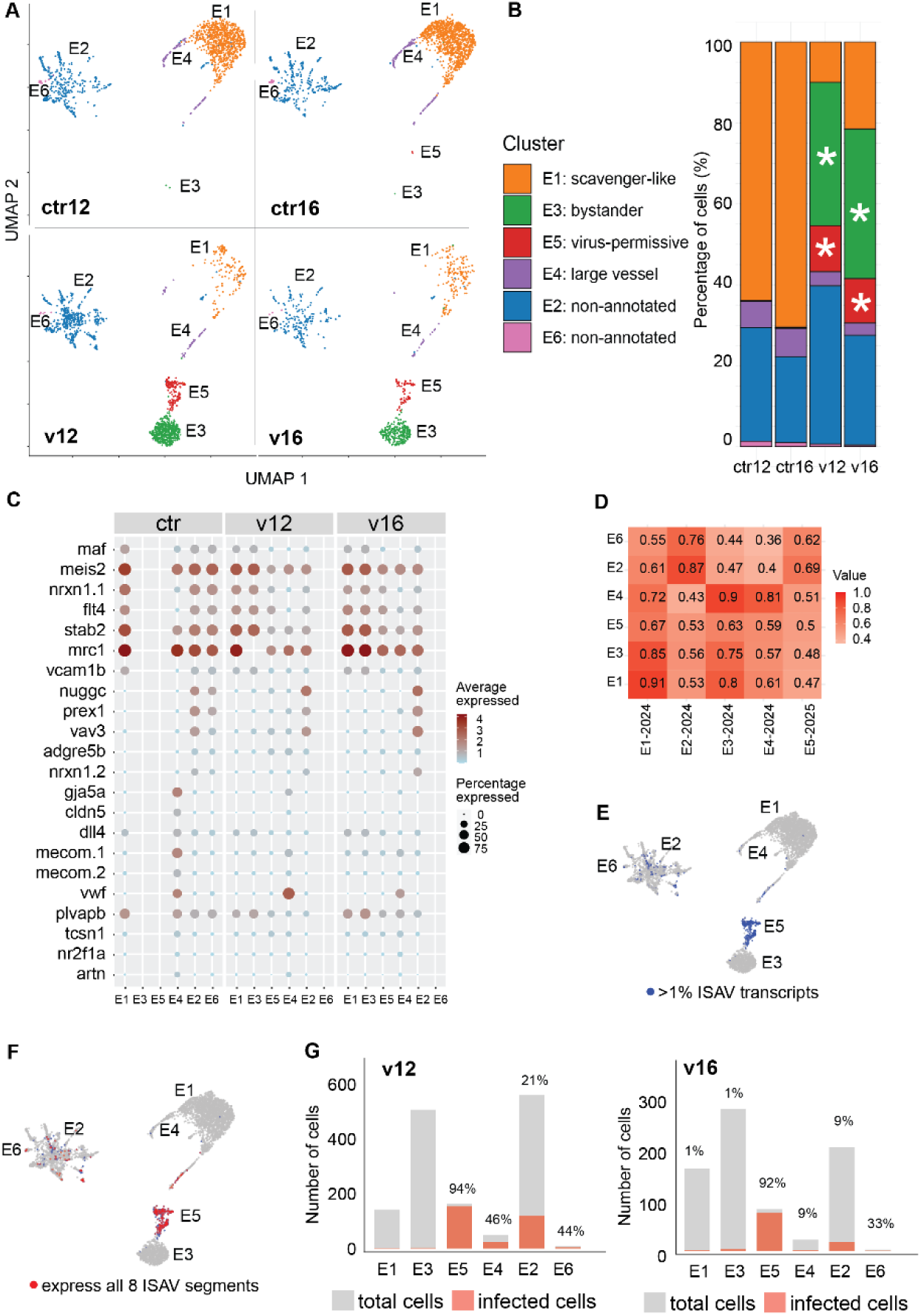
Endothelial cells subpopulations. Endothelial cells represented six subpopulations across sample groups: non-infected fish 12 dpi (ctrl12) and 16 dpi (ctrl16), and infected fish 12 dpi (v12) and 16 dpi (v16). (A) UMAP showing endothelial subpopulations in each group. (B) Barplot showing the percentage of cells in each subpopulation per group. Asterisk highlights those predominantly found in infected groups. (C) Dotplot showing expression of specific marker genes in each subpopulation for each group. In the dotplot, the colour and the dot size reflect the respective average expression and percentage expression for each gene in each subpopulation. (D) Pairwise correlation matrix of average expression of marker genes across endothelial subpopulations with those identified previously in head kidney (18). (E) ISAV gene segment expression across all cell subpopulations and conditions (control and virus-infected, both time points). Cells in blue are considered infected based on at least 1% of transcripts originating from viral RNA. (F) Cells expressing all eight ISAV gene segments simultaneously (shown in red). (E – F) Cells in gray are uninfected. (G) Percentage of infected cells in each subpopulation of infected fish. Ensembl identifiers for all genes shown in the figure are provided in S6 Table.

Two additional subpopulations, E3 and E5, were almost exclusively detected in infected fish (Fig 4B). E3 contained 35% of total endothelial cells in infected fish at 12 dpi and 27% at 16 dpi, while E5 contained 11% of total endothelial cells in infected fish at both 12 and 16 dpi (Fig 4B). When comparing E3 and E5 to the previously identified endothelial cell populations in healthy fish, E3 was closest to the scavenger-like endothelial cells (R = 0.81), while E5 showed limited similarity to previously identified endothelial subtypes (Fig 4C-D). The coinciding emergence of E3 and E5 with a marked reduction in E1 and possibly E4 (Fig 4B) may reflect that a subset of activated cells transitions into a new phenotype in response to infection. Our inference that E3 represents activated scavenger-like endothelial cells is supported by the presentation of E1 and E3 as a single subpopulation in a complementary UMAP that examined the distribution of E1, E3, and E5 from infected fish only (S4 Fig). While the connection of E5 to the other endothelial clusters was more elusive than for E3, the infection-related decline in the E1 population together with the transcriptomic similarity to E3 suggests that at least a proportion of cells in E5 also originated from E1. ISAV-infected cells were primarily located within the infection-typical subpopulation E5 and, to a lesser extent, E2 and E4 (Fig 4E-F). In contrast, cells in E3 expressed very few viral transcripts, suggesting that these cells exhibited a true bystander response.

### Response of bystander endothelial cells during infection

To explore the connection between endothelial cell transcription profiles and infection state, we examined the infection-associated subpopulations in more detail, starting with the bystander endothelial cells. The top 30 marker genes for E3 included *rnf213* (ENSSSAG00000078539), *helz2a* (ENSSSAG00000073332), *interferon-induced 44-like* (ENSSSAG00000038498, ENSSSAG00000107768, ENSSSAG00000045256), *socs1* (ENSSSAG00000118579, ENSSSAG00000104513), the chemokine *scyb7* (ENSSSAG00000106328), and *namptb* (ENSSSAG00000077480), which has a central role in NAD+ biosynthesis (28) (Fig 5A).

**Fig 5.**
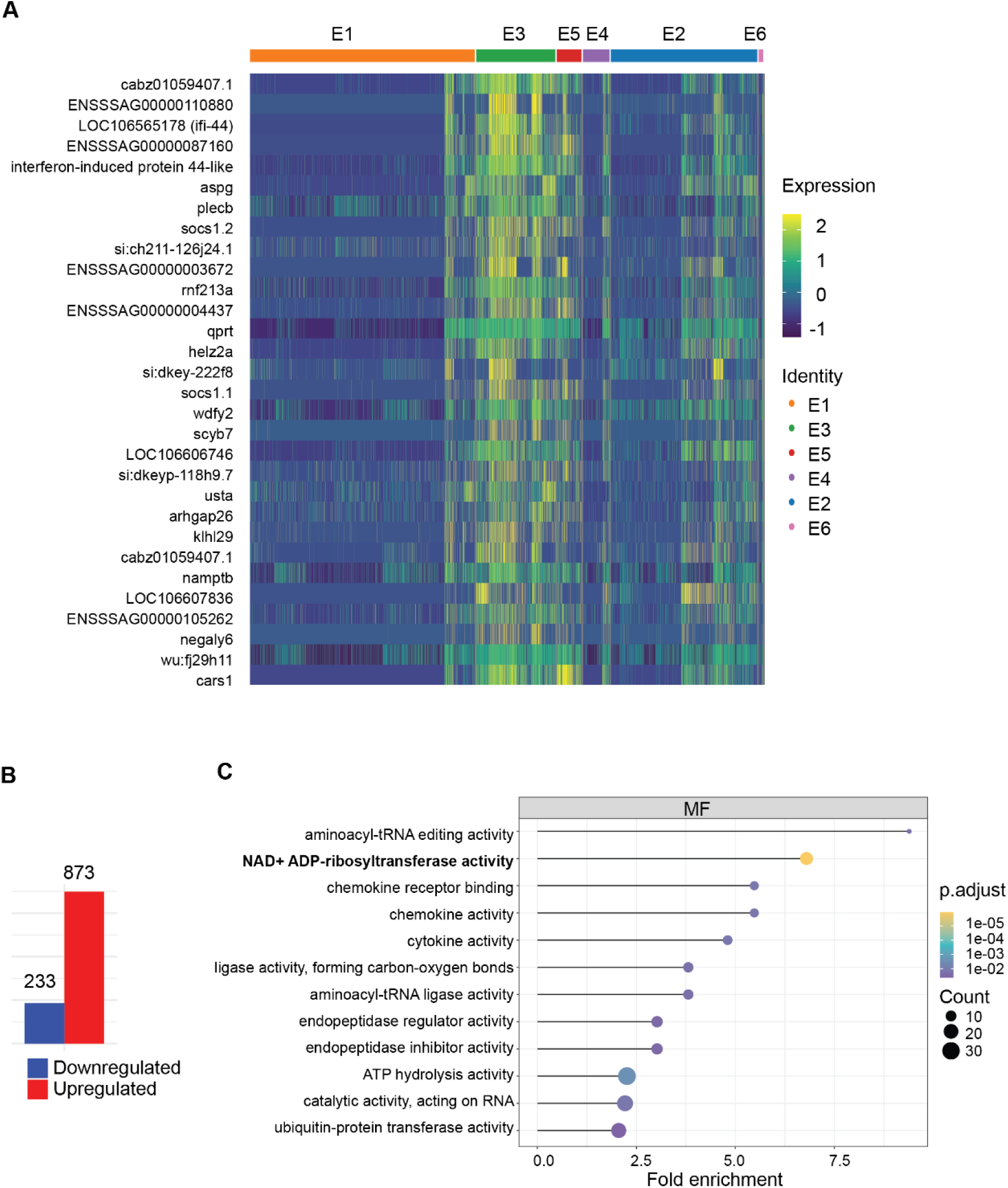
The bystander endothelial response to ISAV infection. (A) Heatmap showing the top 30 marker genes of subcluster E3, annotated as bystander endothelial cells in infected fish. (B) Barplot shows up (red) and down-(blue) regulated genes when comparing E1E3_inf cluster with E1_ctrl. (C) Dotplot showing results of GO analysis of the genes in (B). GO analysis of endothelial response, including both up and downregulated genes using a padj cutoff of 0.05. In the dotplot, the line represents fold enrichment, the colour indicates padj and the dot size the gene count in each term. Ensembl identifiers for all genes shown in the figure are provided in S6 Table.

To further understand the endothelial bystander response to viral infection, we performed a pseudobulk differential expression analysis that compared bystander scavenger-like endothelial cells from infected fish (inf) to scavenger-like endothelial cells from non-infected fish (ctrl). Given the difference in the size of cell clusters in infected (E1_inf = 300, and E3 = 768) and non-infected fish (E1_ctrl = 1893), as well as the transcriptional similarity between E1 and E3 in infected fish (S4 Fig), we merged E1 and E3 from the infected group into one cluster (E1E3_Inf). Overall, 1106 genes were differentially regulated in bystander scavenger-like endothelial cells (E1E3_Inf) of infected fish compared to scavenger-like endothelial cells from the control group, including 873 upregulated and 233 downregulated genes (Fig 5B, S7 Table).

Gene ontology analysis reinforced the observed trends among top marker genes, revealing enrichment of functions related to antiviral and immune responses, such as NAD+ADP-ribosyltransferase activity, cytokine and chemokine activity, and ubiquitin-protein transferase activity, together with those related to protein translation and glycosylation, such as aminoacyl-tRNA ligase and glycosyltransferase activity (Fig 5C, S8 Table).

### Characteristics of directly infected endothelial cells

To investigate the endothelial response to direct infection, we next explored the E5 cluster, detected exclusively in infected fish and with more than 90% of cells classified as virus-infected (Fig 4G). Among the top 30 host markers were genes with assumed roles in immune cell adhesion and interactions, such as *sele* (ENSSSAG00000067000), *alcama* (ENSSSAG00000054868), *cxl10* (ENSSSAG00000061390), and *cd83* (ENSSSAG00000118596) (29–32); tumour suppressors and oncogenes regulating cell cycle, apoptosis, and stress responses, like *urgcp* (ENSSSAG00000096238), *arrdc3a* (ENSSSAG00000007639, ENSSSAG00000071237), *klf6a* (ENSSSAG00000078883), *rassf5* (ENSSSAG00000103465), *dusp1* (ENSSSAG00000037616), and *ier2* (ENSSSAG00000058660); the pluripotent cytokine *tgfb1* (ENSSSAG00000052708); and *serpine1* (ENSSSAG00000068013), a protease with dual antiviral and cytopathic functions (33–39) (Fig 6A).

**Fig 6.**
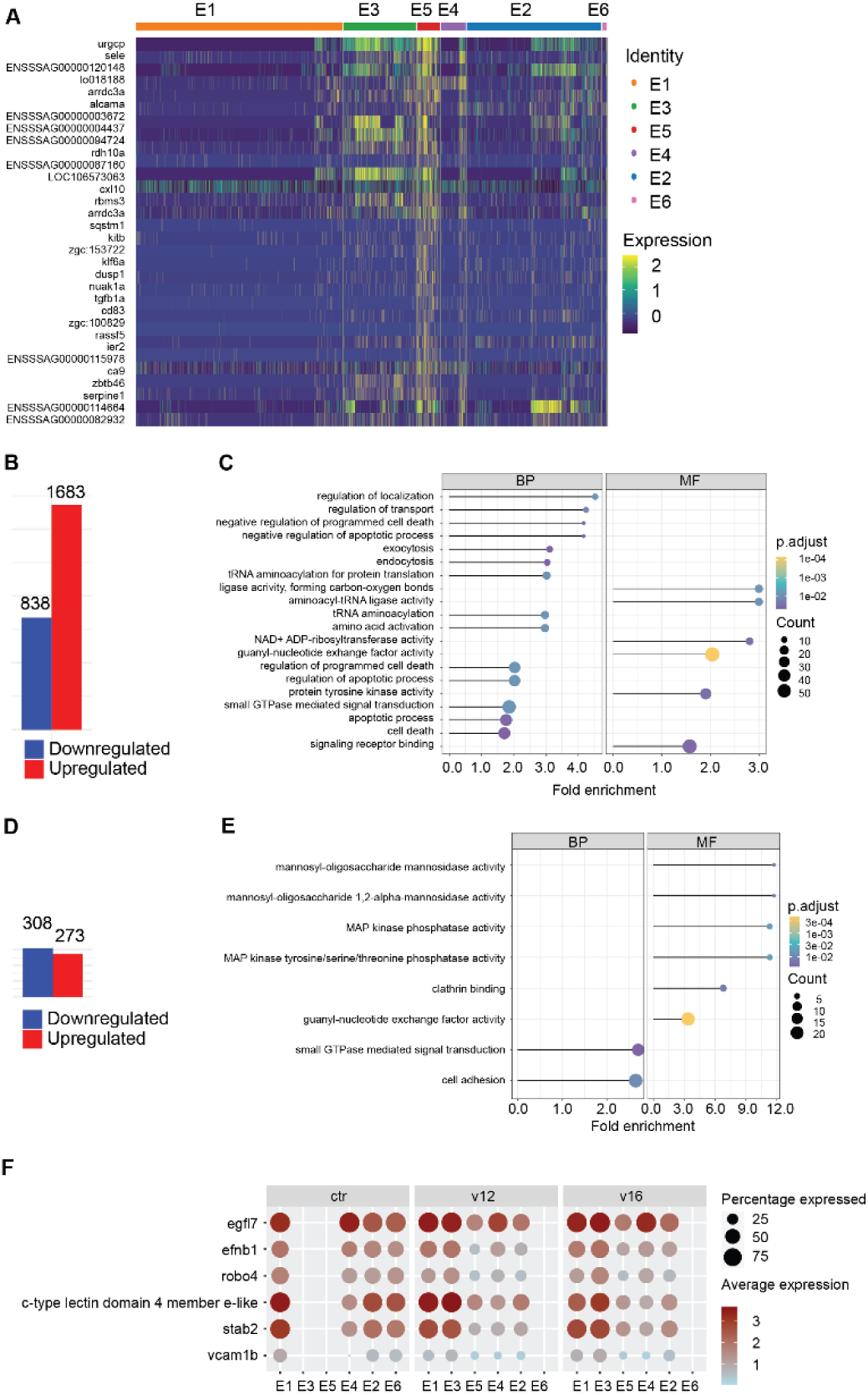
The transcriptomic profile of directly infected endothelial cells. (A) Heatmap showing the top 30 marker genes of E5, representing directly infected endothelial cells in infected fish. (B) Endothelial response to ISAV infection comparing E5 with scavenger-like endothelial cells from non-infected fish (E1_ctrl). Barplot shows up (red) and down-(blue) regulated genes. (C) Dotplot showing results of GO analysis of the genes in (B). (D) Endothelial response to direct infection comparing E5 cluster with bystander-activated endothelial cells in infected fish (E1E3_Inf). Barplot shows up (red) and down-(blue) regulated genes. (E) Dotplot shows results of GO analysis of the genes in (D). (F) Dot plot illustrating loss of general endothelial, as well as specific scavenger-like endothelial markers in directly infected endothelial cells (E5). (C and E) GO analysis of endothelial response, including both up and downregulated genes using a padj cutoff of 0.05. In the dotplot, the line represents fold enrichment, the colour indicates padj and the dot size the gene count in each term. Genes where only Ensembl ID are shown, are non-annotated genes. Ensembl identifiers for all genes shown in the figures are provided in S6 Table.

When comparing the virus-permissive subpopulation (E5) to scavenger-like endothelial cells of non-infected fish (E1_ctrl), 2527 DEGs were identified (838 downregulated, 1683 upregulated) (Fig 6B). This comparison must be interpreted with caution, as the origin of cells in E5 was less clear than E3. Gene ontology analysis confirmed enrichment for terms associated with regulation of apoptosis and programmed cell death along with terms involved in regulation of localization, transport, exocytosis, endocytosis, and protein synthesis (tRNA aminoacylation); small G protein-mediated control of cellular functions (guanyl-nucleotide exchange factor activity and small GTPase-mediated signal transduction); protein tyrosine kinase activity; signaling receptor binding; and NAD+ADP-ribosyltransferase activity (Fig 6C, S8 Table).

A direct comparison of infected (E5) and bystander (E1E3_Inf) endothelial cells within infected fish, identified 581 DEGs, of which 273 were upregulated and 308 downregulated (Fig 6D). Differential enrichment was observed for terms associated with cell adhesion, small G protein signaling (small GTPase signaling transduction, guanyl-nucleotide exchange factor activity), MAP kinase phosphatase activity, clathrin binding, and mannosyl-oligosaccharide mannosidase activity (Fig 6E, S8 Table). Finally, genes associated with endothelial identity (and scavenger-like endothelial identity specifically), were downregulated in E5 compared to E1_ctrl and/or E1E3_Inf, exemplified by *egfl7* (ENSSSAG00000083641), *efnb1* (ENSSSAG00000115237), *robo4* (ENSSSAG00000051277), *tie1* (ENSSSAG00000045503), *c-type lectin domain family 4 member e-like* (ENSSSAG00000120844), *stab1* (ENSSSAG00000067902), and *stab2* (ENSSSAG00000045007) (40) (Fig 6F).

## Discussion

This study presents the first characterization of the transcriptomic response to ISAV infection in individual cell types of Atlantic salmon. It is also the first work to define endothelial transcriptomic profiles during active viral replication of these cells in any species. Our findings provide novel perspectives on the pathogenesis of ISA and factors that may influence Atlantic salmon susceptibility to ISAV infection and disease. In addition, they offer broader insights into the general patterns by which endothelial cells respond to viral infection.

Consistent with previous reports based on ultrastructural, immunohistochemical, and virus binding analyses (5, 41), the vast majority of ISAV RNA was detected in endothelial cells. This aligns with the commonly accepted notion that infection of endothelium is required for ISAV amplification and disease progression. A small fraction of erythrocyte nuclei contained ISAV RNA 12 dpi, in line with our previous observation that these cells, albeit rarely, may express ISAV proteins (6). In contrast, we observed no evidence for ISAV replication in immune cell nuclei, such as mononuclear phagocytes and lymphocytes, strengthening the assumption that these cell types are not permissive to ISAV replication.

The fraction of infected endothelial cells was higher at 12 dpi than 16 dpi, consistent with the observation that ISAV RNA levels in blood increase at 12 dpi but peak at 16 dpi with a subsequent drop (17). Several mechanisms are likely to mediate this decline. First, ISAV particles in blood extensively bind red blood cells, which are removed from the circulation in large numbers (17) probably contributing to immune clearance of the virus. Second, ISAV exhibits strong homologous attachment interference (17, 42), which is likely to limit the infection of new endothelial cells as viraemia progresses. Complementary, previous exposure to viral mimetics inhibits ISAV replication in cultured cells via attachment-independent mechanisms (42). This suggests that cell-intrinsic antiviral response stimulated by soluble signaling molecules and/or viral products can increase the resistance to viral replication as the infection progresses, motivating a detailed examination of the endothelial response to ISAV infection.

Our analysis revealed that many of the endothelial genes regulated by ISAV infection at 12 and 16 dpi were similar and, moreover, overlapped with known interferon-stimulated genes (23). Nevertheless, there was a shift in the overall endothelial response pattern between the two time points. At 12 dpi, when viraemia was increasing and no clinical signs had developed, the response pattern was dominated by pattern recognition (NOD-like) receptor signalling and NAD+ADP-ribosyltransferase activity, with additional enrichment for regulation of programmed cell death, chemokine activity, cytokine activity, and ubiquitin-protein transferase activity. Interestingly, two previous studies observed differential regulation of genes involved in inflammasome activation (related to NOD-like receptor signalling) and ubiquitination between susceptible and resistant fish in early ISAV infection (43, 44). In contrast, NAD+ADP-ribosyltransferase activity, which showed the strongest enrichment both in terms of magnitude and significance, has to our knowledge not been highlighted as a feature of the host response to ISAV infection. In viral infection, NAD+ is used as a source of ADP-ribose, which is transferred to viral proteins in an attempt to inactivate them. In the long run, this may lead to depletion of cellular NAD+ levels (45). Depletion of endothelial NAD+ during oxidative stress and ageing is linked to cytoskeletal disruption and vascular dysfunction (46–48), which may be relevant, considering the pathology of ISA. Enforcing the relevance of vascular dysregulation, several key regulators of vascular functions were enriched at 16 dpi, including apelin signalling, insulin signalling, cellular senescence, protein tyrosine signalling, and semaphorin receptor activity (49–53), together with NAD+ADP-ribosylation and cytokine signalling. As NAD+ levels may be enhanced by dietary supplements (45), our findings advocate for further investigations into the role of NAD+ADP-ribosylation and its effects on cellular NAD+ levels in the context of ISAV pathogenesis.

Exploring the heterogeneity within the endothelial population, we found that infected fish showed a strong reduction in the fraction of endothelial cells with a scavenger-like endothelial profile, which is the most abundant endothelial cell profile in the Atlantic salmon head kidney (18). This reduction was most pronounced 12 dpi, before gross pathology or clinical signs developed, suggesting that the cells had not been lost due to cytopathic effects. More likely, the reduction was caused by extensive reprogramming, shifting scavenger-like endothelial cells to an alternative transcriptional state. Correspondingly, two endothelial subpopulations were exclusively detected in infected fish. One of these subpopulations (E5), contained more than 90% of the directly infected cells at both 12 and 16 dpi. E5 was characterized by a general reduction in expression of endothelial-specific markers, consistent with viral take-over of cellular functions. Moreover, alternative markers emerged, including stress response genes, such as *ier2*, *dusp1*, and *klf6a* (33, 35, 54), and inflammatory mediators that could potentiate cd8+ T cell interactions, such as *sele*, *cxl10*, and *cd83* (29, 30, 32). Enrichment analyses revealed regulation of terms associated with GTPase signalling networks, MAPK signalling, and programmed cell death. These pathways are closely interlinked: Small GTP-ases, activated by guanine nucleotide exchange factors, act as molecular switches with on (GTP-bound) and off (GDP-bound) states. They are upstreams activators of the MAPK pathway that again regulate cell growth, death, and survival (55). Moreover, GTPases control cellular vesicle trafficking, affecting all stages of the viral infectious cycle (56–58). Not surprisingly, different GTPase signalling networks are often hi-jacked by viruses to promote efficient uptake, replication, and release of new viral particles (58–60). This may also affect vascular functions, as GTPase signalling is an important regulators of vascular health and permeability (61–63).

The other infection-typical subpopulation (E3) was, together with the scavenger-like cells, the only subpopulation that did not contain any infected cells. It showed a marked resemblance to the non-activated scavenger-like endothelial cells, likely representing the same cell type. These bystander cells were marked by potentially protective genes, such as *helz2a*, *rnf213*, and *ifi-44* (34, 64, 65). Based on our data, it is not possible to determine if this profile was acquired before encountering infective virus, as a response to soluble mediators, or upon encountering viral components, through pattern-recognition. In any case, the profile is associated with minimal viral replication in an environment where all endothelial cells have most likely been exposed to infection (17), supporting the hypothesis that E3 represents a protective endothelial profile. Understanding endothelial responses that limit viral replication is important, both due to the close connection between ISAV pathogenesis and endothelial replication, and because different endothelial response patterns have been linked to viral disease susceptibility in other species, such as the enhanced susceptibility of black swans to avian influenza (9) or the different susceptibility of sheep and cattle to bluetongue (8). Another marker of this subpopulation was *namptb*, encoding an enzyme involved in NAD+ biosynthesis, potentially countering NAD+ depletion (45).

The relative abundance of E3 and E5 was similar at 12 and 16 dpi, with the proportion of infected cells in E5 remaining high. The decline in infected cells was therefore mainly caused by massive reductions in the proportion of infected cells within the E2 and E4 subpopulations. The reason for this difference is not clear, but one could speculate that E5 represents end-stage infected endothelial cells from all subpopulations, which have lost many of their defining features. Considering the dramatic decline in the scavenger-like endothelial population upon infection, it is not unlikely that a proportion of this population has become infected.

In conclusion, our study has uncovered important information about the Atlantic salmon endothelial response to ISAV infection, strengthening the evidence that ubiquitination and inflammasome activation constitutes a central part of the host response to ISAV and highlighting NAD+ as an interesting and potentially nutrient-sensitive target for further investigation. Moreover, our observations of the temporal changes in the endothelial response to infection, as well as two distinct infection-typical endothelial states, are of general interest and point to possible mechanisms by which endothelial dysfunction could arise as a result of viral infection.

## Materials and methods

### Virus and titer determination

ISAV was propagated in ASK (Atlantic salmon kidney) cells (66) at 15°C, in L-15 medium supplemented with fetal bovine serum (FBS, 5%), L-glutamine (4 mM), and penicillin/streptomycin/amphotericin (1%) or gentamicin (50 µg/mL). All reagents were obtained from Lonza unless otherwise specified. Supernatants were harvested when cytopathic effects were close to complete, 14-28 dpi. Infective titers were determined by inoculating serial dilutions of supernatants in quadruplicate wells of ASK cells cultured in 96-well microtiter plates. Acetone-fixed cells were incubated with IgG_1_ against the ISAV nucleoprotein (P10, Aquatic Diagnostics Ltd, 0.4 µg/mL, 60 min, RT), washed with PBS three times, and incubated with Alexa488-labelled goat anti-mouse IgG (A11001, Thermo Fisher Scientific, 5 µg/mL, 45 min, RT). Titers were calculated using the modified Kärber method, as previously described (67).

### Fish and experimental infection

Infection trial was conducted in agreement with the regulations enforced by the National Animal Research Authority, using protocols pre-approved by the Norwegian Food Safety Authority FOTS (ID: 24382).

Atlantic salmon (AquaGen, Trondheim, Norway) were hatched, reared, and housed at the aquaculture research station: Center for Sustainable Aquaculture (Norwegian University of Life Sciences [NMBU], Ås, Norway). The fish were kept under a 24-hour light photoperiod in circular tanks in a temperature-controlled (14 ± 1°C) freshwater recirculatory aquaculture system. They were fed a standard salmon diet in excess via automatic belt feeders. Before the infection trial, a batch of fish was tested, by qPCR, for infectious salmon anemia virus (organs tested: gills, heart and kidney), salmon pox gill virus (organ tested: gills), infectious pancreatic necrosis virus, piscine rheovirus-1, piscine myocarditis virus, and salmonid alphavirus (organs tested: heart and kidney) by the diagnostic services at the Norwegian Veterinary Institute. All tested fish were found negative for these pathogens.

Altogether, 73 fish with a median body weight of 115 g were included in the infection trial. The experiment was conducted at the NMBU infection aquarium for salmonids in Ås, Norway, which uses a fresh water flow-through aquaculture system maintained at 12°C. After acclimatization, one group (n = 47) was infected by a 2-hour immersion in water containing the highly virulent Norwegian ISAV isolate NO/Glesvaer/2/90 (16), at a concentration of 10^3.75^ TCID_50_/mL, following established protocols (68). For the snRNA-seq study, fish were sampled at 12 (v12) and 16 (v16) days dpi, with three fish sampled per time point. Control non-infected fish, from the same batch, were also sampled (n = 3) at each time point, ctrl12 and ctrl16 respectively. At the time of sampling, fish were euthanized by immersion in an overdose of benzocaine (40 mg/L), weighed, measured, and signs of disease were recorded. Head kidneys were either preserved in formalin for RNAscope in situ hybridization or snap-frozen in liquid nitrogen. Snap-frozen samples were stored in liquid nitrogen until single nuclei isolation.

### snRNA-seq sample preparation

Snap-frozen head kidney samples were shipped to the Roslin Institute Genomics Platform Facility (Scotland), where single nuclei isolation was performed. Nuclear isolation followed a Tween with salts and Tris buffer (TST) based method adapted from (69). Briefly, for each fish approximately 3 mm of flash frozen head kidney tissue was placed in a 6-well tissue culture plate with 1 mL TST buffer. The tissue was minced for 5 minutes on ice using Noyes spring scissors. The homogenate was then filtered through a 40-µm cell strainer, followed by the addition of 1 mL of TST buffer and 3 mL of 1X ST solution was added to achieve a final volume of 5 mL. The sample was centrifuged in a swing bucket centrifuge at 500 g for 5 minutes at 4 °C. The resulting nuclei pellet was re-suspended in 1 ml of PBS 1% BSA, followed by a second filtration through a 40-µm cell strainer. Nuclei were counted manually using a haemocytometer (1:1 Trypan blue) and re-suspended in PBS 1% BSA to an optimal concentration of 1000 nuclei per µl and were subsequently re-counted prior to snRNA-seq library preparation. A detailed list of reagents and catalogue numbers is available in S9 Table.

### cDNA library preparation and sequencing

The snRNA-seq libraries were prepared at the Roslin Institute Genomics Platform Facility using the Chromium Next GEM Single Cell 3’ Reagent Kits v3.1 (10x Genomics) following the manufacturer’s protocol targeting 6000 nuclei per sample. Briefly: Nuclei, barcoded RNA capture beads and reagents for reverse transcription were loaded onto the chip and placed into a Chromium controller where nuclei were individually partitioned into nanodroplets with barcoded RNA capture beads. The nuclei were subsequently lysed and mRNA reverse transcribed with an added barcode, droplets were broken and pooled cDNA was amplified and quality controlled (QC) using an Agilent Tapestation HSD5000 assay (Agilent Technologies). cDNA passing QC was then prepared for sequencing by fragmentation, end-repair, A-tailing, ligation, and indexing PCR. The libraries were QC using an Agilient Tapestation HSD5000 assay, before submitting for sequencing. Sequencing was performed at the Wellcome Trust Clinical Research facility, Western General Hospital, Edinburgh, on the NextSeq 2000 platform (Illumina Inc.) using NextSeq 1000/2000 P3 Reagents (200 cycles) v3 Kit, 1.2 B paired-end reads following the manufacturer’s protocol.

### Data analysis

The current Atlantic salmon reference genome (Ssal_v3.1; GCA_905237065.2) was downloaded from Ensembl version 106 (70).

The mitochondrial assembly and annotations from the previous reference (ICSASG_v2; GCA_000233375.4) were appended to this reference. STARsolo (2.7.10a) (71) was used to map the raw reads (fastq files) to this genome, demultiplex and error correct cell barcodes, and quantify per-cell gene expression. The ‘STAR’ command was used with the following parameter changes: –-outSAMtype BAM SortedByCoordinate, –-soloType CB_UMI_Simple, –-clipAdapterType CellRanger4, –-soloFeatures GeneFull_Ex50pAS, –-soloUMIdedup 1MM_CR, –-outFilterMatchNmin 40, –-outFilterScoreMin 40, –-soloBarcodeReadLength 0, –-soloUMIfiltering MultiGeneUMI_CR, –-soloCellFilter EmptyDrops_CR, –-soloMultiMappers EM. Summary statistics for mapping rates are provided in S10 Table.

The raw count matrices were processed using Cellbender (0.3.0) (72) to identify and eliminate ambient RNA signals and to perform cell calling. Optimal parameters for expected nuclei numbers and the total droplets to include were determined using ranked barcode plots, with the number of epochs set to 200. Quality control for each dataset was conducted separately using Rstudio (73) with the package Seurat (4.4.0) (74). Nuclei with fewer than 150 genes or 150 UMIs, as well as those with more than 10% of UMIs delivered from mitochondrial genes, were removed from downstream snRNA-seq analysis. After excluding poor-quality nuclei, all 12 samples were merged. Quality control parameters were re-analyzed on the merged data, leading to further improvement to exclude potentially compromised nuclei, including those suspected of being damaged or doublets. Specifically, we adjusted the mitochondria threshold to 5% and added a ribosomal protein gene threshold of 2.5%. We also excluded nuclei with gene counts higher than 7000 and UMI counts higher than 16000 to remove potential doublets. The “findDoubletClusters” command from the R package scDblFinder (1.12.0) (75) was also used to identified and remove homotypic doublets – doublets formed from transcriptionally similar cells.

The merged data was normalized using “lognormalization” with standard Seurat parameters for visualization, clustering, and UMAP generation. Integration was performed using Harmony (76) to correct batch effect between samples. Gene annotations were retrieved from the Ensembl database for Atlantic salmon. Due to a large number of unnamed genes in the salmon genome annotation, biomart (77) was used to identify orthologues in other species, including salmonids, northern pike, zebrafish, medaka, chicken, mouse, and human. For genes without annotations, the gene name of the most closely related orthologue was assigned (18, 78). Ensembl gene IDs are included in the text and supplementary tables to differentiate the many paralogues retained from two successive whole genome duplication events in teleost and salmonid evolution (79–81).

We performed automated annotation of the main cell populations using specific markers for each Atlantic salmon head kidney cell lineage, as defined from our previous work (18). Cluster annotation started by performing high-resolution clustering (resolution 8 in the “findClusters” Seurat command) and annotating each cluster using the following criteria: mean absolute expression of markers > 0.1; at least 50% of cells expressing all marker genes; and the mean expression > 0.5 standard deviations above the mean expression for the dataset.

Initially unassigned clusters were manually inspected and given appropriate identities. Clusters displaying two distinct identities were flagged as potential doublets. Additionally, scDblFinder (75) was used again in the merged data to detect doublet clusters, and those that aligned with our automated annotation were excluded.

Pseudobulk analysis was performed (82) to identify differentially expressed genes between conditions (control vs. virus-infected) in each major cell population, analyzing the two time points separately (ctrl12 vs. v12 and ctrl16 vs. v16). Raw counts were extracted and aggregated at the sample level, and differentially expressed genes were identified using DESeq2 (83). ISAV segment genes were excluded from the analysis, as they were only present in infected groups and could skew the results. Genes were considered differentially expressed if they showed a Log2FC > 1 for upregulated genes or < –1 for downregulated genes, with an padj < 0.05.

To understand the heterogeneity within endothelial cell, we created a subset of all endothelial cells and applied the same analysis steps as for the whole dataset to identify subpopulations within this cell type.

To assess the correlation between endothelial subpopulations from this study and our previous work (18), we used the R function “cor.” The analysis was based on the average expression levels of all marker genes, calculated for pairwise combinations across all endothelial subpopulations in both studies. Codes used for running the analysis and plotting are publicly available on GitHub repository at github.com/NorwegianVeterinaryInstitute/Salmon_HighLowISAV. The data discussed in this publication have been deposited in NCBI’s Gene Expression Omnibus and are accessible through GEO Series accession number (to be assigned).

### In situ hybridization

Head kidney samples were collected from three fish at 12 dpi. The fish used for RNAscope were not the same individuals as those used for snRNA-seq. Tissues were fixed in 10% formalin for at least 24 hours, dehydrated, embedded in paraffin, and thin sections (4 µm) were placed on slides. An RNAscope™ 2.5 LS probe for V-Salmon-Isavirus (Cat. No. 847528) was used to target ISAV segment 8 (HQ259678) (84). Hybridization was performed using the RNAscope LS Red automated kit (ACD Bio, Cat. No. 322750) on a Leica BOND RXm platform. As a positive control, the RNAscope™ 2.5 LS Probe for Ssa-ppib (Cat. No. 494428) was used to detect *Salmo salar* peptidylprolyl isomerase B (cyclophilin B). A probe targeting dihydrodipicolinate reductase (dapB) from *Bacillus subtilis* (ACD Bio, Cat. No. 312038) was used as negative control.

## Acknowledgements

We would like to thank Eirill Aager-Wick and Hetron Mweemba Munang’andu (the NMBU infection aquarium for salmonids, Ås, Norway) for administrative and technical support during infection trial; Simon Chioma Weli for support during the infection trial; Ricardo Tavares Benicio and Bjørn-Reidar Hansen (the aquaculture research station, NMBU, Ås, Norway) for providing fish and technical assistance; Rose Ruiz Daniels for assistance during nuclei isolation; and the bioinformaticians at the Norwegian Veterinary Institute for technical support. Computations resources used for preliminary analysis were provided by Sigma2 – the National Infrastructure for High-Performance Computing and Data Storage in Norway. We acknowledge the contribution of the Roslin Institute Genomics Platform Core Facility to this study.

## Supporting information captions

**S1 Fig. Marker genes and percentage of cell types in Atlantic salmon head kidney**. (A) Violin plot showing the expression level of marker genes for each cell population. (B) Barplot showing the percentage of cells in each cluster for each fish: non-infected fish 12 dpi (ctrl12) and 16 dpi (ctrl16), and infected fish 12 dpi (v12) and 16 dpi (v16) (n = 3). Proportion of infected cells in (C) endothelial cells and (D) erythrocytes across infected fish samples. Percentages indicate the fraction of infected cells relative to the total cell count in that sample. Thromb – thrombocytes, NK-L – natural killer-like cells, MPs – mononuclear phagocytes, Mesen – mesenchymal cells, Inter – interrenal cells, HSCs – haematopoietic stem cells, Granu – granulocytes, Eryt – erythrocytes, and Endo – endothelial cells. Ensembl identifiers for all genes shown in the figures are provided in S6 Table.

**S2 Fig. ISAV detection by in situ hybridization**. In situ hybridization of formalin-fixed, paraffin-embedded tissue sections revealed ISAV RNA expression in endothelial cells lining thin-walled sinusoids and larger vessels in the head kidney (A-B), heart (C-D), and liver (E-F). ISAV segment 8 was targeted using the RNAscope™ 2.5 LS Probe for V-Salmon-Isavirus. The figures are from fish in the v12 group (12 dpi).

**S3 Fig. Heterogeneity of the E4 endothelial cluster**. (A) UMAP visualization of merged samples highlighting subpopulation 4 of endothelial cells, which splits into two distinct clusters. (B) Feature plots showing the expression of key marker genes within subpopulation E4. Colors indicate normalized expression levels: gray denotes no expression, while increasing intensities of purple represent higher log-transformed expression values. Displayed genes include *mecom* (ENSSSAG00000048049), *gja5a* (ENSSSAG00000042461), *vwf* (ENSSSAG00000101168), and *lama5* (ENSSSAG00000108070).

**S4 Fig. Specific endothelial subclusters in infected fish**. (A) UMAP visualization of endothelial cell subclusters E1, E3, and E5, including cells from infected fish only.

**S1 Table. Complete list of markers from single nucleus RNA sequencing analysis**. The table shows the log2 average expression of each gene in each cell population. Only genes expressed in at least 20% of the cells (min.pct = 0.2) and with a log2 fold change ≥ 0.5 were included. Bonferroni correction (threshold p < 0.05) was applied to filter out non-significant markers. A numerical suffix was added to distinguish gene names that appeared multiple times on our annotation list with unique Ensembl ids.

**S2 Table. Number of infected cells in different cell populations**. Cell were considered infected when at least 1% of its transcripts originated from viral RNA.

**S3 Table. Differentially expressed genes from pseudobulk analysis**. Differentially expressed genes identified in pseudobulk analysis at 12 dpi and 16 dpi, analyzed separately. Genes were classified as differentially expressed if Log2FC > 1 (upregulated) or Log2FC < –1 (downregulated), with an adjusted p-value < 0.05.

**S4 Table. KEGG and Gene Ontology (GO) enrichment analyses of the endothelial response.** KEGG and GO enrichment analyses were performed on endothelial cells isolated at 12 and 16 days post-infection (dpi), referred to as Endo12 and Endo16, respectively. Terms and pathways were considered significantly enriched if the adjusted p-value was less than 0.05. Fold enrichment was calculated as the ratio of the observed gene count to the expected gene count for each pathway (KEGG) or term (GO).

**S5 Table. Endothelial subpopulations markers**. Table displaying the log2 average expression of each gene across different cell populations. Only genes expressed in at least 20% of the cells (min.pct = 0.2) and with a log2 fold change ≥ 0.5 were included. Bonferroni correction (threshold p < 0.05) was applied to filter out non-significant markers. A numerical suffix was added to gene names that appeared multiple times in the annotation list, corresponding to unique Ensembl IDs.

**S6 Table. List of marker genes included in figures**.

**S7 Table. Differentially expressed genes from pseudobulk analysis comparing Endothelial cell subpopulations.** The table presents differentially expressed genes identified in pseudobulk analysis of endothelial cell subpopulations. Genes were classified as differentially expressed if Log2FC > 1 (upregulated) or Log2FC < –1 (downregulated), with an adjusted p-value < 0.05.

**S8 Table. KEGG and Gene Ontology (GO) enrichment analyses of the endothelial response.** E1E3_Inf – bystander-activated endothelial cells in infected fish; E5 – directed infected endothelial cells; E1_ctrl – scavenger-like endothelial cells from non-infected fish. Terms and pathways were considered significantly enriched if the adjusted p-value was less than 0.05. Fold enrichment was calculated as the ratio of the observed gene count to the expected gene count in each term (GO).

**S9 Table. List of reagents used for nuclei isolation. S10 Table. Mapping stats**

