## Supplementary figures and images for "Virus-inclusive single-nucleus RNA sequencing reveals two distinct endothelial response patterns in infectious salmon anaemia"

### Supplemental figure 1

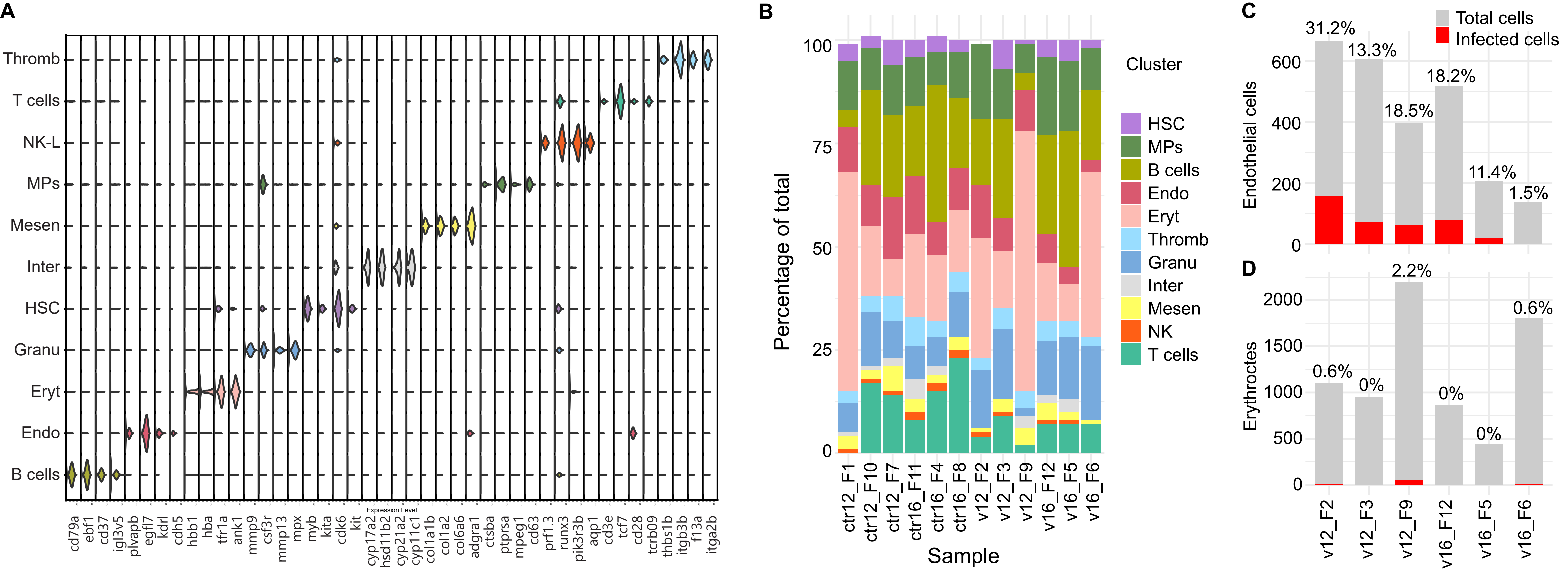

### Supplemental figure 2

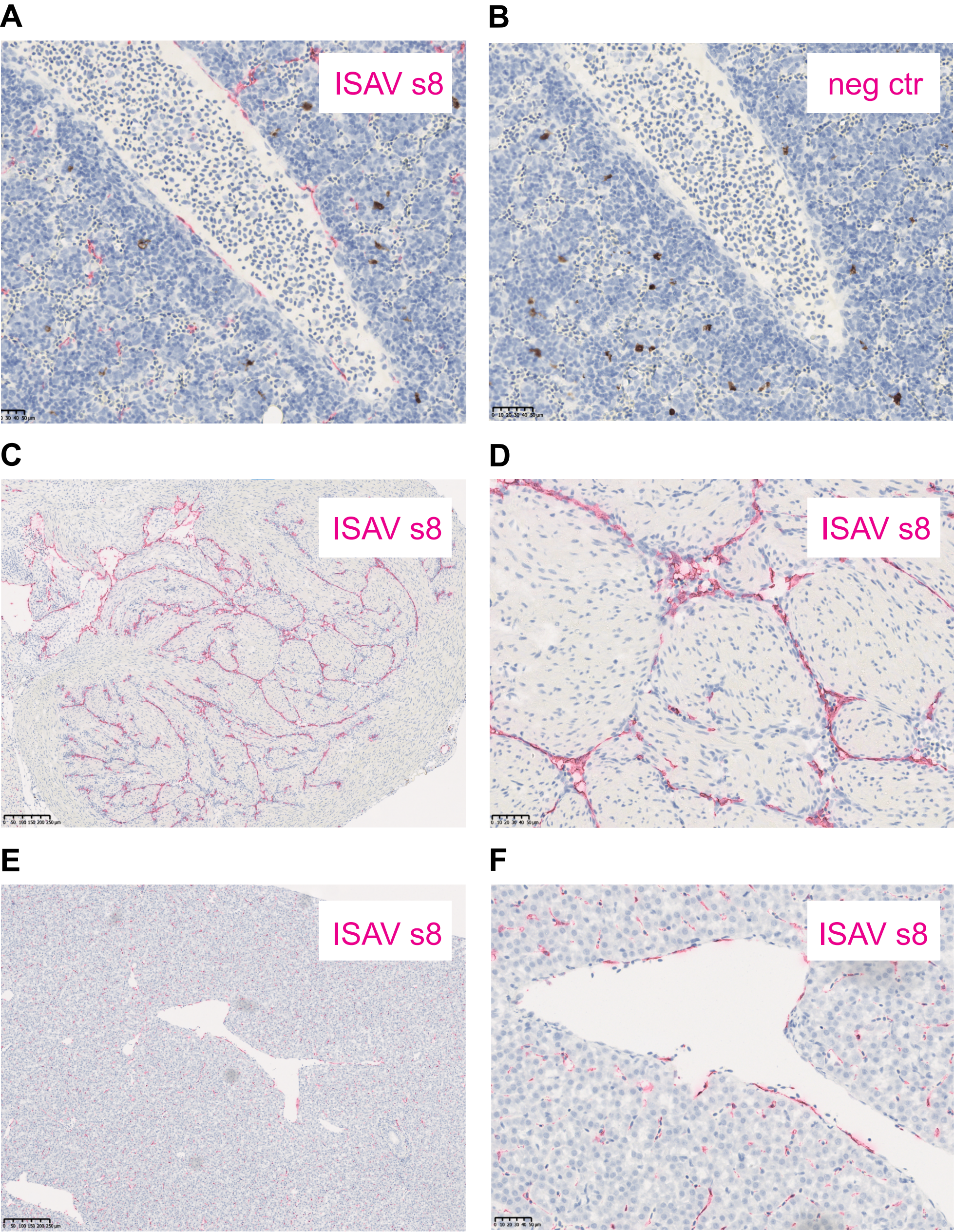

### Supplemental figure 3

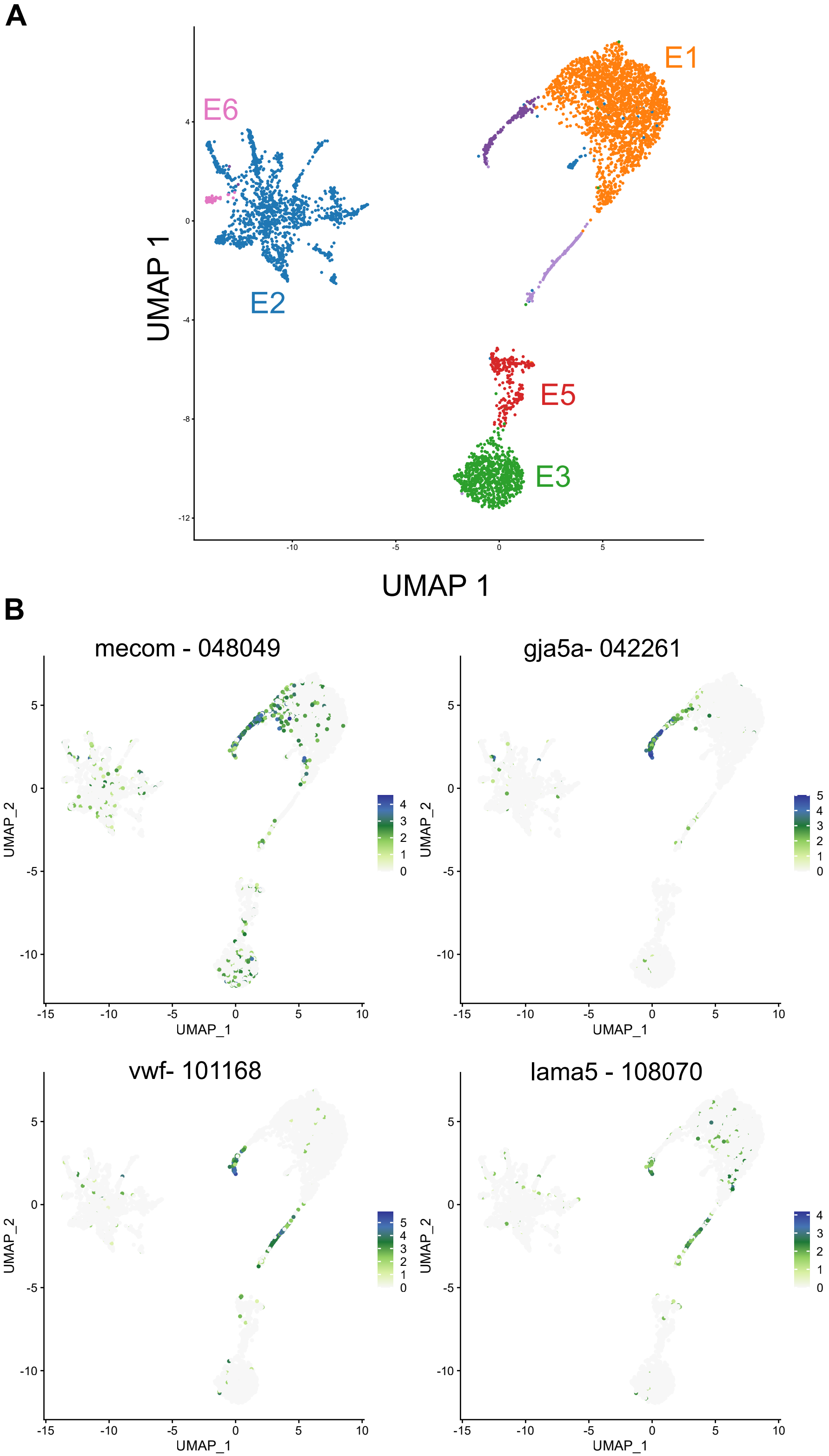

### Supplemental figure 4

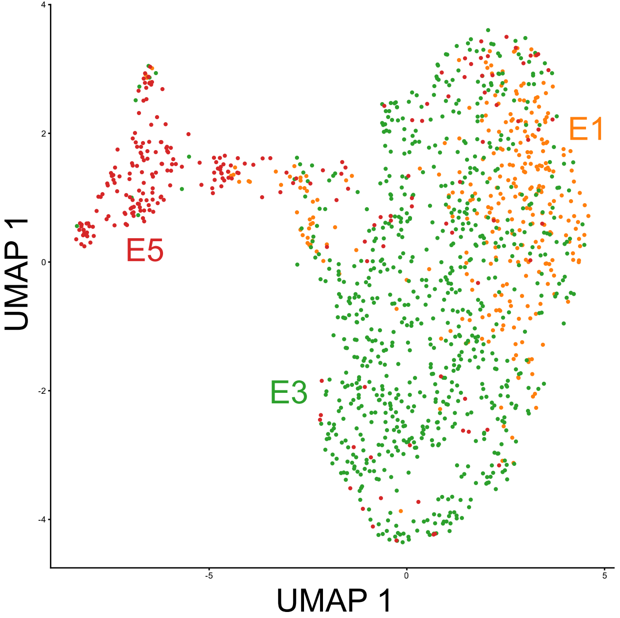
